# Profiling the Expression of Transportome Genes in cancer: A systematic approach

**DOI:** 10.1101/2023.07.18.549498

**Authors:** Luca Visentin, Giorgia Scarpellino, Luca Munaron, Federico Alessandro Ruffinatti

## Abstract

The transportome, the *-omic* layer encompassing all Ion Channels and Transporters (ICTs), is crucial for cell physiology. It is therefore reasonable to hypothesize a role of the transportome in disease, and in particular in cancer. Here, we present the Membrane Transport Protein DataBase (MTP-DB), a database collecting information on ICTs, and a pipeline that takes expression data and the MTP-DB as input to produce a broad overview of transportome dysregulation in cancer. The MTP-DB may prove useful for the study of the transportome in general, and the pipeline may be used to study the transportome in other diseases. Both tools are open source and can be found on GitHub at TCP-Lab/mtp-db and TCP-Lab/transportome_profiler, under permissive licenses. We detect that the transportome is dysregulated in cancer, and that dysregulation patterns are shared among different cancer types. It is still unclear how these patterns are linked to cancer patho-physiology.

## 1 Introduction

In each kingdom of life, the diffusion of substances between the intracellular and extracellular environments is essential for cell survival. Apart the free diffusion of small lipophilic molecules, these exchanges are mediated by transmembrane proteins of various nature, which we refer to by the term *transmembrane transport proteins*, while the*-omic* layer that comprises all of them is referred to as the *transportome*.

The coordinated action of these proteins regulates a large number of physiological functions, such as membrane potential, nutrient absorption, waste product removal, cellular signaling, regulation of intra- and extracellular pH, and more.

While the expression of these proteins and their proper targeting are necessary pre-requisites for membrane transport, they are generally not sufficient for the fulfillment of the overall function. In fact, the establishment of Transport Functional Units (TFUs), which are capable of performing specific tasks, often requires the assembly of multiple protein subunits to form homo-or heteromultimers, or even long-range interactions among different transmembrane transport proteins. For instance, any secondary active transporter cannot function properly in the absence of a pump that creates a chemical gradient to be dissipated. Another example is given by the communication between *STIM* calcium sensors and *Orai* channels, responsible for the calcium release-activated calcium currents.

The cataloging and characterization of TFUs can be a complex task, especially due to current limitations in proteomics research and the still low-throughput nature of the biophysical techniques for functional assays. Currently, according to a widely accepted approximation, the individual TFU is identified with the gene transcript of one or more subunits it consists of, assuming transcriptional levels to be directly proportional to the abundance and activity of the TFUs they encode. Although this approximation may have many limitations (poor correlation between transcripts and proteins, lack of functional assessment, unclear role of auxiliary subunits, …), it serves as a practical approach until more precise methods for TFU quantification and characterization become available.

According to the schema in Figure 1, transport proteins can be broadly classified into two main categories: *pores* and *transporters*.

**Figure 1.**
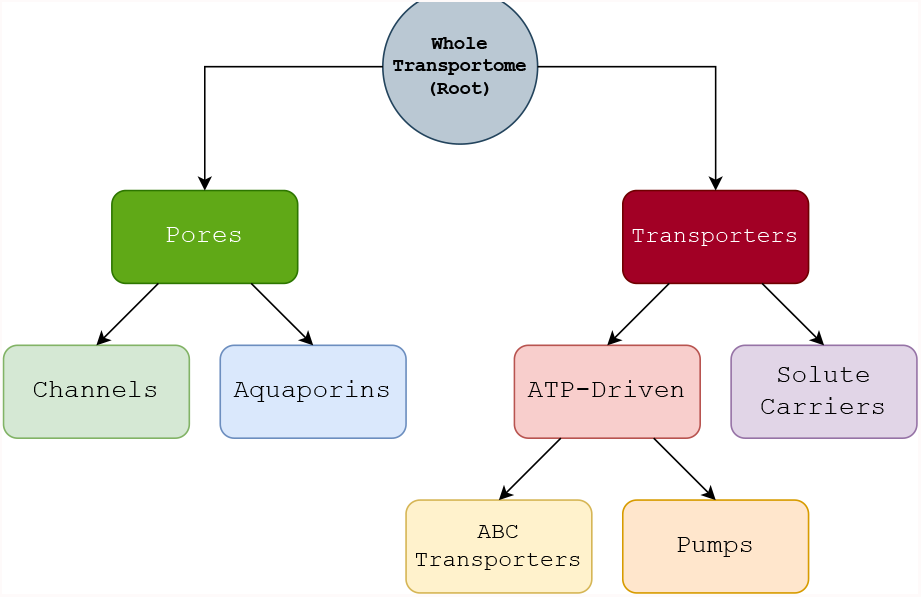
Tree of the principal classes of ICTs.

- **Pores:** water-filled pores that allow the facilitated passage of molecules through the membrane. These may be additionally subdivided into *ion channels* proper and *aquaporins*, pores that mostly allow the passage of water.
- **Transporters:** proteins or protein complexes that, embedded in the membrane, allow the passage of molecules upon conformational changes. These may be furthermore divided into those that require ATP hydrolysis to function and those that do not, known as *ATP-driven (or primary active) transporters* and *SoLute Carriers (SLCs)*, respectively. A common distinction within the ATP-driven transporters is given by *ATP-Binding Cassette (ABC) transporters* and *pumps*, where ABC transporters feature the conserved ABC domain, while pumps do not.

Overall, the phrase “Ion Channels and Transporters (ICTs)” is now commonly used to refer to the set of all the transportome gene products that fall into one of these macro-categories.

Cancer cells exhibit fundamental differences from their healthy counterparts, especially in their relationship with the extracellular environment. Dramatic metabolic shifts are also observed in cancer. Both of these aspects probably involve an alteration in the transportome, acting as an “adapter”^1^ for cancer cells with the tumor micro-environment, while at the same time ensuring the exchange of nutrients and metabolites capable to sustain the altered metabolism. By the above approximation, this transportome dysregulation may be reflected in the expression levels of the genes themselves, making it interesting to explore the transcriptional profile of cancer cells.

One commonly employed approach to accomplish this task is by measuring gene expression in both healthy and diseased samples. This is followed by Differential Expression Analysis (DEA) and enrichment analysis of the resulting gene list using ontologies—such as the Gene Ontology (GO)—and hypothesis tests like hypergeometric test or Fisher’s exact test. Finally, the list of all the significantly enriched terms needs to be screened *a posteriori* to search for some transportome-related terms. The effectiveness of this process heavily relies on both the statistical power reached by the DEA and the careful curation and organization of the ontology database.

A similar—but somehow “reversed”—approach is to generate a limited number of gene sets of interest *a priori*, which meaningfully group together genes belonging to, for instance, similar TFUs. Then a Gene Set Enrichment Analysis (GSEA) can be performed to test them against the data and see if the function or the gene family they represent is dysregulated or not. This second option has a few advantages:

- The weighted GSEA method can take into account the magnitude of differential expression of the genes in the list (i.e., the effect size), and not their mere presence or absence.
- The tested gene lists may be arbitrary and not necessarily based on ontologies. For example, they may be manually curated, specifically crafted for a purpose (such as a list of genes involved in a specific function or pathway of interest), or generated by other methods.
- Given a set of features, it is possible to systematically generate all the gene sets that may be meaningfully conceived, and test them all against the data.

The present work aims at profiling the expression levels of transportome genes in the context of human cancer. To do this, we collected information on these genes (such as their complete list, which molecule(s) they transport, their gating mechanism(s), their functional class, etc.) and used it to systematically arrange them into meaningful Transporter Gene Sets (TGSs). After sorting all the protein-coding genes found in cancer cells based on their differential expression with respect to healthy cells, we ran a pre-ranked GSEA on these ordered lists to obtain enrichment scores for every TGS. We therefore obtained the “dysregulation status” of most functional facets of the transportome in 19 different cancer tissue types.

We provide an open-source, documented, and reproducible Python package, Daedalus (github.com/TCP-Lab/MTP-DB) that retrieves transportome-related data from various databases and compiles it in a local `.sqlite` database. In parallel, we also provide pre-compiled database files as periodic releases.

A `make`-driven and docker-containerized pipeline, named “transportome profiler”, (github.com/TCP-Lab/transportome_profiler) is also available. It takes gene expression data and the aforementioned database to generate gene sets, sorts genes based on their differential expression, and runs GSEA. This pipeline was designed with modularity and reproducibility in mind, so that it would be easily adaptable on other datasets and databases.

## 2 Materials and Methods

In this section we broadly cover the implementation of the pipeline, with particular regard to the generation of the TGSs and the related GSEA.

### 2.1 The Membrane Transport Protein Database

Even if the GO knowledgebase [1, 2] represents an invaluable source of meaningful and readily available gene sets, we decided not to rely solely on the manually curated and automatically generated gene lists of the GO. Instead, we preferred to explore all the possible TGSs that can originate by the transporter-related features of interest in a more unbiased and systematic way.

To improve the ease of use and portability, we compiled all the data we needed to generate the TGSs in a SQLite database and named it Membrane Transport Protein DataBase (MTP-DB). To populate the MTP-DB, we sourced information from a variety of online databases. In particular, we extracted data from the Human Gene Nomenclature Committee (HGNC) database [3], the International Union of basic and clinical PHARmacology (IUPHAR) “Guide to Pharmacology” database [4], Ensembl [5] through the use of BioMart [6], the Transporter Classification DataBase (TCDB) [7], the SLC Tables website [8], the Catalogue Of Somatic Mutations In Cancer (COSMIC) [9] database, and the GO [1, 2].

The specific data downloaded from each database and the methods used to retrieve them can be seen in Table 1. A Python package nicknamed *Daedalus* was created to download, parse, and save this information, recreating the MTP-DB *on the fly* at every execution. We opted for this generation method instead of a manually updated, static database file for a few reasons: a dynamically regenerated database will always be up-to-date in respect of the upstream database changes, it does not require particular hosting capabilities to be accessed even in the far future (as the users can simply download and run the source code to obtain a copy), and it provides a clear, transparent, and traceable way to see how the source data is obtained, parsed, and stored (by examining the source code).

**Table 1:**
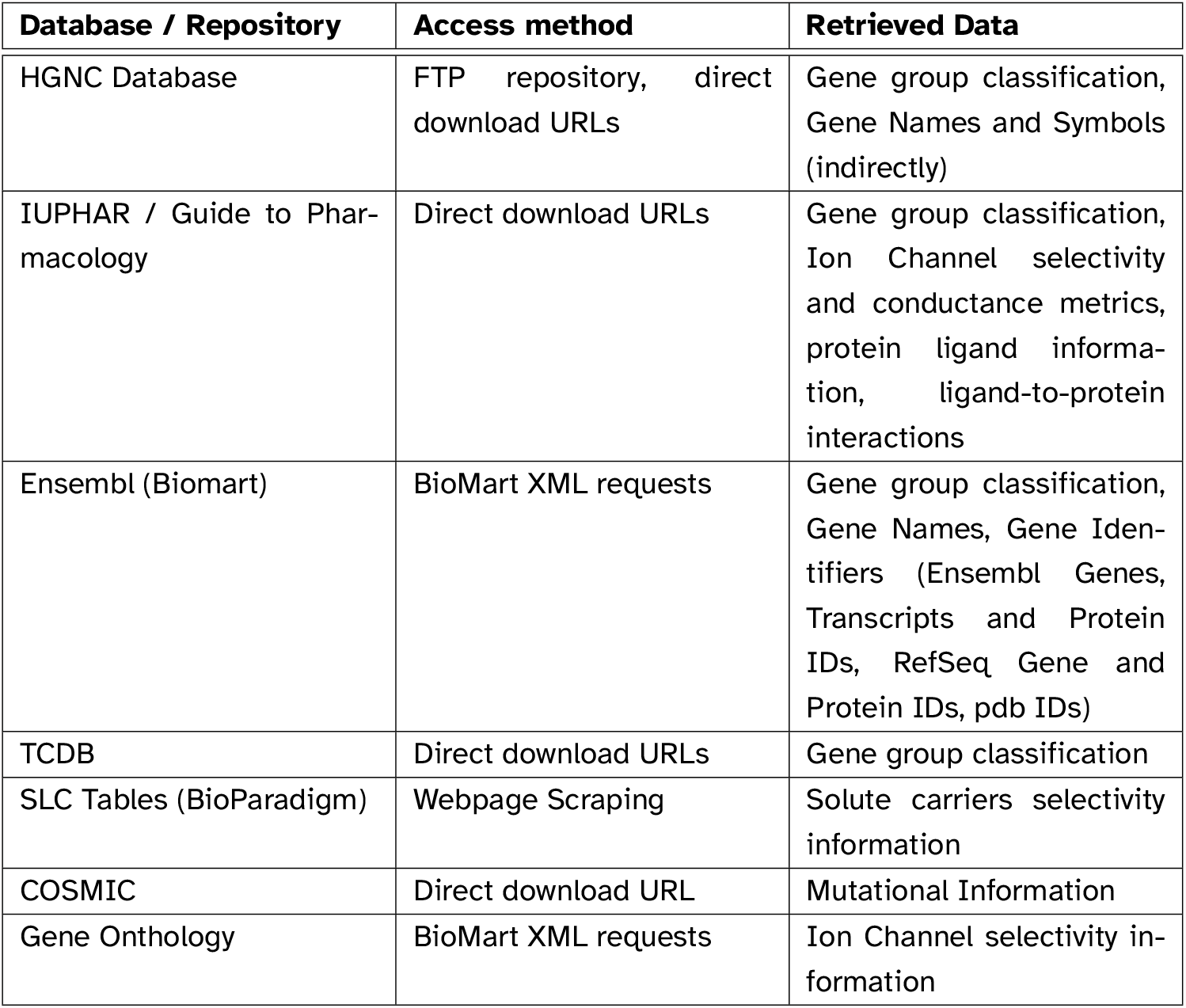
List of accessed databases, methods of access, and the use of the downloaded data in the Membrane Transport Protein DataBase.

| **Database / Repository** | **Access method** | **Retrieved Data** |
| --- | --- | --- |
| HGNC Database | FTP repository, direct download URLs | Gene group classification, Gene Names and Symbols (indirectly) |
| IUPHAR / Guide to Pharmacology | Direct download URLs | Gene group classification, Ion Channel selectivity and conductance metrics, protein ligand information, ligand-to-protein interactions |
| Ensembl (Biomart) | BioMart XML requests | Gene group classification, Gene Names, Gene Identifiers (Ensembl Genes, Transcripts and Protein IDs, RefSeq Gene and Protein IDs, pdb IDs) |
| TCDB | Direct download URLs | Gene group classification |
| SLC Tables (BioParadigm) | Webpage Scraping | Solute carriers selectivity information |
| COSMIC | Direct download URL | Mutational Information |
| Gene Ontology | BioMart XML requests | Ion Channel selectivity information |

It was interesting to notice how information from the above databases was, and still is at the time of writing, flawed in several minor ways. For instance, the GABA_A_ receptors in the brain are ionotropic receptors notoriously permeable to chloride ions, but the HGNC database only classifies them as *ligand gated ion channels* and not as *chloride channels*. Additionally, the IUPHAR does not contain any permeability information for any of the *GABR\** genes (conductance and permeability being properties of the receptor *protein complex* rather than the single subunit gene). This makes it impossible for Daedalus to classify them as permeable to chloride ions. It is also known that GABA_A_ receptors exhibit a certain permeability to HCO_3_^-^, generally in between 0.2 and 0.4 that of the chloride ions [10], but not even this feature is reflected in the above databases. A similar argument applies to nicotinic acetylcholine receptors (nAChRs) and related *CHRN\** subunits. Moreover, when classifying channels by *gating mechanism*, HGNC makes no distinction between *gating* proper (in the senses of activating stimulus) and *modulation*. As a result, some voltage-modulated channels, such as the TRP superfamily of cation channels are pooled with pure voltage-activated channels such as K_v_, Na_v_, and Ca_v_, which may not be desirable from a physiologist’s point of view. A final example is how some ion channels (including many beta subunits of potassium voltage-gated and calcium-activated channels, as well as the mitochondrial calcium uniporter family of proteins) were missing from the ion channel lists provided by the IUPHAR and the HGNC. All of these defects were manually addressed in the MTP-DB through post-build hooks automatically applied by Daedalus during each build process. Notably, even such “terminal patches” can be easily inspected in detail by the reader in order to get an idea of the degree of manual curation within the MTP-DB. If needed, post-build hooks can be turned off to obtain a version of the database free of manual annotations.

To support a more varied search strategy, especially when querying data for SLCs, we implemented a sort of internal thesaurus, adding synonyms or more general terms to the database besides the already-present information on the carried molecule in all relevant tables. For example, we inserted the more general values of “cation” and “anion” next to each particular ion species; we added more general synonyms such as “carbohydrate” for mannose and glucose; “amino acid” for all basic amino acids; and so on. The thesaurus is also consulted to correct any parsing errors made by Daedalus, and manually addresses some imperfections in the source data (e.g., different codes for the same amino acid are all converted to the three-letter code). The thesaurus is contained in a single, human-readable comma-separated-values file, available in the remote repository.

The MTP-DB will be an effective way to stimulate further research in the Transportome by providing a centralized and human- and machine-friendly place to obtain a lot of relevant information on Transportome genes. The pre-compiled MTP-DB and the Daedalus package are open-source and available for free on GitHub at www.github.com/TCP-Lab/MTP-DB, and we encourage users to inspect and propose improvements.

At every development milestone, a published version of the database (in the form of the generated SQLite file) will be released on GitHub, for reproducibility purposes. The latest release can be downloaded directly from this URL in `gzipped` format. Additionally, note that the data blob downloaded and then used by Daedalus to generate the database is locally saved as a Python Pickle file, and is the only file that the package needs to regenerate the database. This can be useful to exactly reproduce the database in the future even after the remote databases have updated. The database is versioned using the CalVer specification, with representation `Major.YY.0W[_MINOR][-TAG]`.

At the time of writing, the latest version of the database is `1.23.26-beta`, featuring a total of 932 ICT gene entries (356 ion channel, 14 aquaporins, 47 ABC transporters, 83 pumps, and 432 SLCs).

The MTP-DB is, however, still partially incomplete. We manually curated a list of 151 calcium-permeable ion channels and found that, although most of them were correctly annotated as such, 68 (45%) were not. Of these, 30 were annotated as “cation” permeable, but without the more specific term for calcium. In that case, much of the inconsistency stemmed from the different interpretation of the term “calcium channel” given by the list curators. HGNC, for example, reserves this term for only those ion channels *selectively* permeable to Ca^2+^. On the contrary, our list included all calcium-permeable channels, even those non-selective for that ion species. Nevertheless, some well-known calcium-selective ion channels (including *PIEZO1, PIEZO2, P2RX7*, and some others) were not correctly annotated in any of the sources consulted by Daedalus. We addressed these missing annotations, but we cannot however be sure that the same phenomenon does not occur for other ion channels, or even more generally in other transporters. For this reason we set up the GitHub repository with a collaborative approach in mind, following Open Science practices. This would allow researchers that notice an issue with the annotations in the database to participate in the creation of a more useful resource.

A docker image per every release, representing the environment in which the database was rebuilt in, is available on Docker Hub at cmalabscience/mtpdb. Please note, however, that Daedalus does not directly depend on any environment variables.

An overview of the database schema, with information on the type of data in every table, can be seen in Table 2. The actual database schema file is present in the GitHub repository at this URL.

**Table 2:** Brief description of the data contained in all the tables present in the MTP-DB.

| Table name | Description |
| --- | --- |
| gene_ids | Genome-wide gene identifiers from Ensembl (“ENSGs”) and their versions. |
| transcript_ids | Transcript identifiers from Ensembl (“ENSTs”), and their version, associated with gene identifiers. |
| mrna_refseq | RefSeq transcript identifiers and their ENST counterparts. |
| protein_ids | Protein identifiers from Ensembl (“ENSPs”) associated with their respective ENSTs, PDB accession codes and RefSeq protein IDs. |
| gene_names | ENSGs associated with their respective HUGO gene IDs, symbols and names. |
| iuphar_targets | Every gene considered as a “target” by the IUPHAR, with relevant names, IDs and family information. |
| iuphar_ligands | Every ligand considered by the IUPHAR, as well as its type, name, ENSG (if relevant), pubchem SIDs and CIDs, and information on drug status. |
| iuphar_interaction | Every interaction between a IUPHAR-recognized ligand and human protein target, with relevant information. |
| tcdb_ids | The entirety of the TCDB linked with the respective ENSPs, split by type, subtype, family, subfamily and superfamily. |
| tcdb_types | Names of the types in the TCDB. |
| tcdb_families | Names of the families in the TCDB. |
| cosmic_genes | Information regarding the presence of the gene in the COSMIC database, as well as its hallmark status. |
| channels | All genes catalogued as “channels” by IUPHAR or the HGNC, with information on their gating mechanism, carried solute and relative and absolute conductance. Information from GO regarding permeability information is also included. |
| aquaporins | All aquaporin genes as well as their expression tissue. |
| solute_carriers | All genes catalogued as as “solute carriers” by IUPHAR or the HGNC, as well as their carried solute, rate of transport, stoichiometry, port type, direction, and more. |
| pumps | All active transporters (excluding ABC transporters), correlated with their carried solute, rate, transport direction, net transported charge, and more. |
| ABC_transporters | All ABC transporter genes, correlated with their carried solute, rate, direction, and more. |

### 2.2 Generation of Transporter Gene Lists

We created a Python script to query the database file produced by Daedalus and generate the TGSs. We structured the TGSs as a tree in which each node represents a separate TGS. All transportome genes are present in the root node, while subsets of these gene sets are considered in every child node.

To divide the genes into meaningful TGSs, we seeded the algorithm to always produce a manually defined tree—or “core” tree—reflecting the classification schema already shown in Figure 1. This subdivision purposefully follows also the internal schema of the database, since most nodes represent a table or a combination of whole tables. The algorithm then retrieves all relevant information to each node of the core tree and finds all the possible sub-classes based on the available data, finally appending each sub-class to the tree as a child node. This approach is then repeated for each added child, recursively creating smaller and smaller child gene sets.

In practice, for each core node, the database is queried to obtain the relevant columns of data to the starting node, possibly joining multiple tables. For instance, for the “ATP-Driven” core node, the algorithm will consider data from both the `ABC_transporters` and the `pumps` tables. Once the data is retrieved from the database, for each available column the algorithm generates gene groups of unique genes with the same shared column value, appending them to the parent node. The algorithm then considers a subset of the original data, based on the column value used to generate the child gene set, and attempts to create more by considering all other columns in turn. This approach essentially enumerates all possible gene sets that could be meaningfully derived from the annotated data.

This iterative process generates a very large number of gene sets (with version 1.23.25-beta of the database, this number was ∼ 6000). For this reason, we implemented a pruning procedure on the resulting tree, which prunes away leaf nodes that are highly similar with other nodes in the tree.

We measured node similarity by the Sørensen–Dice coefficient κ as defined below:

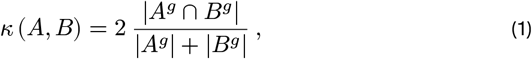

where *A* and *B* are nodes in the tree *T*, associated to the sets of genes *A*^*g*^ and *B*^*g*^, respectively. κ is a value ranging from 0 to 1, where 1 means perfectly congruent nodes, and 0 meaning completely different nodes. Given a tree *T*, we calculated the similarity index for each node *A* as

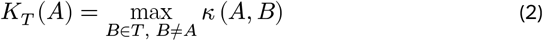

If *K*_*T*_ (*A*) is greater than a user-defined threshold value *η* (generally close to 1), the node *A* is deemed redundant and pruned. After all leaves are considered, if the algorithm pruned at least one leaf, it is executed again. This process returns a tree in which all leaf nodes are sufficiently different from each other. On the contrary, internal nodes are always preserved, even in the presence of high similarity scores between them, since they cannot be eliminated without the removal of meaningful leaf nodes.

The acute reader might notice that, as κ is a symmetric measure, this process removes leaves in an order-dependent way. Indeed, if κ (*A, B*) > *η*, the pruning of *A* or *B* depends on which is considered first. This was taken into account, and the leaf nodes are sorted based on their depth (the number of edges to follow to return to the root node) before the pruning cycle begins. This allows the usage of two pruning strategies: bottom-up or top-down. In the top-down approach, higher leaves (farther from the root) are pruned first, as the nodes are sorted in descending order. The opposite happens in the bottom-up approach, where leaves closer to the root are pruned first, in favor of distant leaves.

Do note that although κ is symmetric, similarity among sets is not a transitive relation (except for κ = 1), so the two pruning methods do not necessarily generate trees with an identical number of nodes. This is not surprising, as three nodes *A, B*, and *C*, where κ(*A, B*) > *η*, κ(*A, C*) > *η*, but κ(*B, C*) < *η* will be pruned differently if they are considered in one order (e.g., *A → B → C*, resulting in {*B, C*}) or the opposite (e.g., *C → B → A*, resulting in {*A*}).

For efficiency purposes, the algorithm does not consider data columns that contain too many missing values (deeming them not meaningful to split on, with the default being > 50% emptiness), does not generate gene sets that are smaller than a threshold (by default 10 genes) and does not consider for further subsetting gene sets smaller than a threshold (by default 40).

The specifications for the core tree, the SQL calls used to retrieve the data for each node in the core tree, the threshold for κ, the pruning direction, and other settings are all easily configurable as command-line arguments. This makes this approach easily adaptable to future versions of the MTP-DB and even other kinds of SQLite and more generally SQL-based databases provided by other researchers. For our run, we used the default parameter values, while κ was set to 0.9.

Once generated, the tree is translated to a set of files in folders on disk. The folders are named with the names of the nodes, and the `all.txt` files in them contain the Ensembl Gene IDs of the genes in the specific TGS. This structure was chosen to easily allow the reconstruction of the tree by any other programs, simply by following the tree-like file system paths.

### 2.3 GSEA analysis

Raw expression data (in the form of un-normalized RNA-Seq counts) and related meta-data to accomplish this analysis were downloaded from the University of California Santa Cruz’s Xena platform [11], selecting the `TCGA TARGET GTEx` study (although TARGET samples were omitted from the analysis). This data have the benefit of being processed by the same analysis pipeline, eliminating technical noise coming from the pipeline itself.

To run GSEA, we needed to pair The Cancer Genome Atlas (TCGA) tumor expression data with their Genotype-Tissue Expression (GTEx) “healthy” counterparts and then perform DEA on each cohort separately with the DESeq2 R package. This proved to be not so straightforward. The Xena platform provides a fused version of the GTEx official metadata file plus the TCGA metadata for ease of use. However, it is unclear how to meaningfully pair such large databases, as methodology, possible batch effects due to sample handling, sample preprocessing (such as microdissection), and the intrinsic non-healthy nature of GTEx samples can cause confounding in the results.

To pair the healthy and tumor data together, we therefore followed the macroscopic grouping provided by the metadata files in the Xena platform. To subset the very large expression matrix file based on the metadata, we implemented a Python package called *metasplit*, which makes use of the low level Rust-compiled `xsv` package to speed up the computation [12]. The specific calls used to subset the expression matrix and therefore match the tumor and healthy samples are available in the GitHub repository at this url. As an overview, we compared tumor samples with their healthy counterparts from the same tissue or organ of origin.

This comparison is purposefully generic. As we lack specific information on the actual location of the biopsied tissue, its specific microdissection status, and potentially other experimental biases or variables at play, we deemed to be safer not to compare different tumor types of the same organ by themselves, but combine them into a more general category.

The fitted DESeq2 model had only the tumor status variable and the intercept. This resulted in an independent Wald-statistic metric for each gene in each comparison. We used the Wald-statistic values as weights to run a weighted, pre-ranked GSEA analysis with the fGSEA R package [13] using the previously generated TGSs of interest. We ran the analysis with both actual values for the statistic and absolute ones, in order to also detect possible symmetric dysregulations. Thus we obtained one significance table for each tumor-healthy cohort, with enrichment information for each input gene set. Finally, we used this data to generate a global enrichment matrix with gene sets on the rows, the cohorts on the columns, and the Normalized Enrichment Score (NES) as values.

## 3. Results

The enrichment matrix generated by the pipeline can be seen in Figure 2. The matrix provides a summarized overview of the over-or under-expression of each TGS across all cohorts. As each leaf node can be considered meaningfully different from any other node, the matrix represents an eagle’s-eye-view of the dysregulation in the different cancer types.

**Figure 2.**
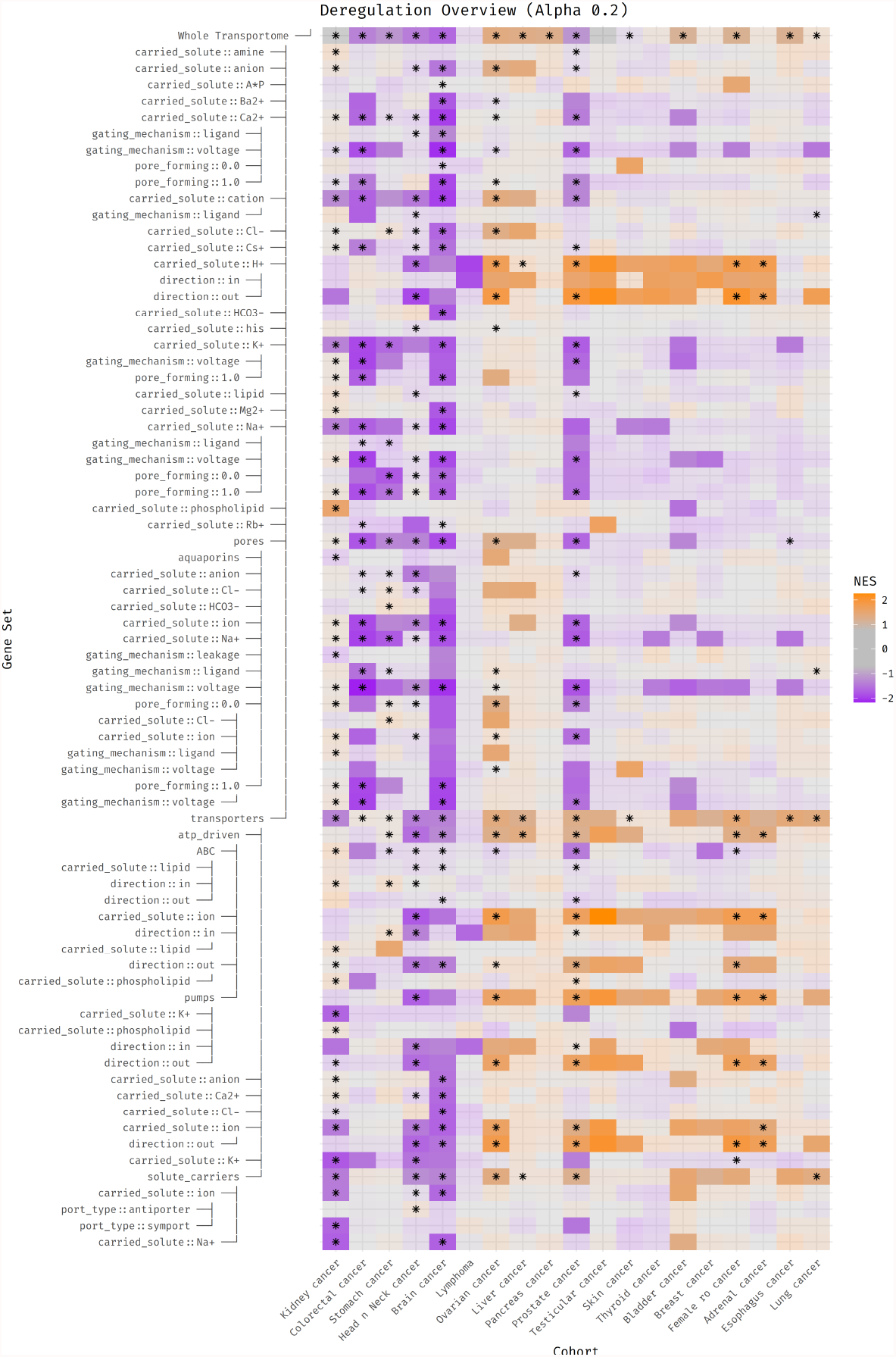
Results of GSEA about transportome dysregulation presented as a heatmap with the NES values for each TGS of interest (as rows) and the tested cohorts (as columns). Columns are clustered hierarchically. Opaque boxes are statistically significant (as reported by adjusted *p*-values from GSEA) to an alpha threshold of 0.20, while semi-transparent ones are not. Coordinates with a dot are significant to the same alpha threshold when considering absolute Wald-statistic values.

In particular, some tumor types closely cluster with various signatures:

- Kidney, colorectal, stomach, head and neck, and brain cancer show generalized dysregulation in many categories, and more specifically downregulation.
- Brain cancer shows downregulation or dysregulation in most categories.
- Pancreatic, esophageal, lung cancer and lymphoma show little to no deregulation, with the exception of a strong downregulation of proton ICTs.
- Prostate, testicular, skin, thyroid, bladder, female reproductive organs, adrenal gland, and breast cancers share a general upregulation of proton ICTs.
- Kidney cancer shows downregulation in a few categories, but shows significant dysregulation when looking at absolute metric values across almost all gene sets.
- The only cancer that shows a difference in expression of aquaporins is ovarian cancer.

Looking at the heatmap row-wise, we are able to appreciate other features:

- With the exception of lymphoma and ovarian cancer, most cancer types show either no change or downregulation of pores.
- Transporters are generally dysregulated. In most cancer types they are upregulated, and seem downregulated in only Kidney, Head and Neck and Brain cancer.
- Not many cancers (just three, colorectal, prostate, and breast) show dysregulation in ABC transporters, but more show absolute dysregulation in the same category.

In conclusion, transportome genes are not uniformly dysregulated in cancer, but rather in a cancer-specific way. However, functional considerations are limited by the fact that the gene sets are not completely mutually exclusive, and that they may not necessarily be functionally homogeneous.

Any implications that these overview has in the broader context of tumor physiology are still to be determined, but it is clear that the transportome may be a key player in cancer biology.

## 4. Other Statements

### 4.1 Author Contributions

Conceptualization, L. V., L. M. and F. A. R; Methodology, L. V. and F. A. R; Software, L. V. and F. A. R; Validation, F. A. R; Formal analysis, L. V., F. A. R. and G. S.; Data Management,

L. V. and G. S.; Writing, L. V., F. A. R and G. S.; Visualization, L. V.; Supervision, L. M.; Project Administration, L. V and L. M.; Funding acquisition, L. M.

### 4.2 Data and Code Availability

Combined TCGA and GTEX data and metadata is available from the Xena platform at this url and this url. The MTP-DB is available on GitHub at TCP-Lab/MTP-DB. The full analysis code as well as `LaTeX` code to generate the author manuscript is available on GitHub at TCP-Lab/transportome_profiler.

## Footnotes

1 Literally in the sense of *mediator of the adaptation* towards the extracellular environment.

